# Dietary variations drive divergent phenotypic, transcriptomic, and metatranscriptomic profiles in *Biomphalaria glabrata*, a schistosomiasis vector snail

**DOI:** 10.1101/2025.09.29.679075

**Authors:** Damilare O. Famakinde, Ciaran Lonergan, Duncan Wells, Geoffrey N. Gobert, Paul McVeigh

## Abstract

**Background:** The freshwater snail *Biomphalaria glabrata* is an important natural vector for the human parasitic trematode *Schistosoma mansoni,* which causes schistosomiasis. In the laboratory, *B. glabrata* are routinely maintained on simple lettuce diets. We aimed to explore and compare the impact of alternative diets on snail performance, global gene expression, and microbiome.

**Methods:** Snails were raised in groups on fresh lettuce (FL), fish food (FF) and artificial snail gel (SG) diet for eight weeks, while measuring dietary impacts on growth, survival, and fecundity. RNA sequencing (RNA-Seq) was performed to correlate dietary phenotypes with changes in the snail transcriptome and associated microbial metatranscriptome.

**Results:** Relative to FL, FF and SG diets markedly enhanced growth, survival, and fecundity, with FF generating the highest fecundity rate. RNA-Seq identified 21,887 nutritionally modulated genes in the snail transcriptome. Fish food (FF) and SG diets drove upregulation of genes associated with antimicrobial immunity, growth, and reproduction, while elevated expression of genes linked to xenobiotic metabolism and oxidative stress was observed in FL-fed snails. Metatranscriptomic analysis identified 104 microbial classes, with a total of twenty-three classes significantly enriched in FF and SG snails, including short-chain fatty acid-producing and nutrient-cycling bacteria. Significant correlation (r = 0.63, *p* = 0.001) linked differentially expressed genes with enriched microbial taxa, highlighting the impact of diet on key snail health and performance metrics.

**Conclusions:** This work is the first nutritranscriptomic analysis of laboratory-bred *B. glabrata*. We describe key insights into the diet–phenotype–transcriptome–microbiome axis, which will inform dietary precision and optimisation for laboratory culture of *B. glabrata*. These data also highlight fundamental aspects of snail biology which could be exploited for molecular snail control approaches.

## Background

*Biomphalaria glabrata* is a neotropical planorbid freshwater snail that transmits *Schistosoma mansoni*, a parasitic blood-dwelling fluke that causes hepato-intestinal schistosomiasis in humans and other mammalian hosts [1, 2]. Schistosome miracidia undergo asexual reproduction within snail tissue, developing through two successive sporocyst generations, eventually releasing cercarial larvae into fresh water to infect the definitive hosts. Mature diecious adult schistosomes subsequently reproduce sexually within the definitive mammalian host, yielding eggs containing further miracidia which hatch when in contact with fresh water [1].

Snail culture is an essential aspect of laboratory maintenance of the schistosome life cycle. The practice of maintaining *Biomphalaria* snails in captivity can be traced back to the early 1900s when various intermediate snail hosts of schistosomes were first identified [3]. Since then, colonies of *B. glabrata* have been maintained in laboratories worldwide to support the life cycle of *S. mansoni* [4] and echinostomatid flukes [5], and as a model for exploring new potentials for snail vector control [6]. *Biomphalaria glabrata* is also a useful model for ecotoxicological risk assessments [7] and even for studying complex human diseases [8].

Maintenance of *B. glabrata* populations under experimental conditions depends on a range of factors, amongst which diet is crucial. However, the choice of diet varies from one laboratory to another, ranging from single natural diets to complex formulated ones. Natural diets used in *B. glabrata* husbandry include green lettuce (*Lactuca sativa*), *Spirogyra*, cyanobacteria and microalgae [9, 10]. Formulated snail diets used may be either semisynthetic such as alginate snail food, and fish food [6, 11] or crude organics such as multi-agal mixtures [9]. While these more complex diets are useful for specific purposes, literature and informal conversations with the research community show that fresh, wilted or dried lettuce is the staple single food source in many laboratories. This reflects its low cost, accessibility, and apparent palatability for the snails.

In freshwater snails that transmit trematodes, including *Biomphalaria* spp., *Bulinus truncatus*, *Galba truncatula* and *Lymnaea stagnalis*, a range of phenotypic traits are influenced by diet. These include growth, fecundity, mortality, snail/fluke association [12–15], tolerance to toxicants [16, 17], and various biochemical indices [18, 19]. The most compelling datasets on *Biomphalaria* snails hypothesize that low-nutrient and high-nutrient diets are directly proportional to growth, survival and egg production, as well as cercarial output in schistosome-infected snails [13, 20–22]. While these dietary effects have clear implications for snail culture in controlled settings and for schistosome transmission dynamics in the wild, we do not understand the molecular mechanisms behind their phenotypic impacts.

Diet, as a key regulator of metabolic processes, influences physiology via complex networks, including regulation of gene expression, either directly by activation of transcription factors, or indirectly through epigenetic methylation of DNA and modification of histones, impacts on regulatory non-coding RNAs, or modification of microbial populations [23, 24]. Following completion of the Human Genome Project [25], the term ‘nutrigenomics’ (nutritional genomics) was introduced and describes all forms of nutrient– genome interactions [26], including ‘nutritional transcriptomics’, which focuses on the effects of nutrient intake on gene expression [27]. Nutritional transcriptomics, or nutritranscriptomics, has been studied in a range of human and non-human species [28, 29], and this approach is the focus in this study.

Our study compared the impact of lettuce and relatively simple artificial food sources on key parameters relevant to laboratory maintenance of *B. glabrata*, including survival, growth and reproductive output. We also investigated the nutritional transcriptomic basis of these phenotypes through analysis of snail gene expression as well as metatranscriptomic analysis of the microbial communities associated with snail tissue in distinct diet groups. Our data identifies key factors in dietary optimisation for *B. glabrata* laboratory culture, and highlights key aspects of snail biology representing potential targets for molecular snail control.

## Methods

### Experimental snails

Uninfected Naval Medical Research Institute (NMRI) strain *B. glabrata* snails were used in this study, originally provided in 2020 by Dr Gabriel Rinaldi from the Wellcome Sanger Institute, Hinxton, UK. Snails were maintained in aquaria at Queen’s University Belfast, following standard protocols by the Biomedical Research Institute (BRI) Schistosomiasis Resource Center (Rockville, MD, USA) (https://www.afbr-bri.org/schistosomiasis/standard-operating-procedures/).

### Feeding assays and phenotypic data analysis

Initially, seventy viable size-matched hatchlings (~ 1 mm in size) were isolated from a nursery aquarium and separated into seven feeding groups, each containing ten hatchlings. Each group was maintained on a separate single or mixed diet as follows: (i) fresh green butterhead lettuce (*Lactuca sativa var. capitata*) (FL, purchased commercially), (ii) dried lettuce (DL; prepared from FL leaves, oven-dried at 100 □ for about 10h until completely dehydrated and then crushed into small fragments), (iii) wilted lettuce (WL; wilted mechanically by hand rubbing), (iv) snail gel food (SG; prepared as described previously [30], containing barley grass powder (Sevenhills Wholefoods, UK), wheatgerm (Your Health Store, UK), tropical fish food flakes (Pets at Home, UK), powdered milk (Your Health Store, UK), and sodium alginate (Special Ingredients, UK), in 8:2:2:1:2 ratio). Final diet groups included (v) Love Fish tropical fish food flakes (FF; Pets at Home, UK), (vi) WL supplemented with SG (WL+SG), and (vii) WL supplemented with FF (WL+FF).

Snails were kept in transparent, aerated 1L plastic containers filled with 500 mL of diluted 1x artificial river water (10x ARW: 5L H_2_O, 2.78g CaCl_2_, 6.15g MgSO_4_.7H_2_O, 2.1g NAHCO_3_, 215mg K_2_SO_4_ and 250µL FeCl_3_.6H_2_O, pH 7.0), at 25 □ constant room temperature and under artificial 12:12 light/dark cycle for eight weeks. Snails were fed *ad libitum* three times per week. Prior to each feeding, the containers were cleaned and ARW was replaced. In the mixed-diet groups, the diets were swapped at one-week intervals. Based on performance across the observed phenotypes, three monospecific diets (FL, FF, and SG) were selected for nutritional transcriptomics analyses. In this assay, each dietary feeding was initiated with fifteen size-matched hatchlings, which were later sampled for RNA sequencing at eight weeks post-feeding. By the end of all feeding experiments, each dietary regimen has been repeated at least twice.

During the feeding period, growth, reproductive output and mortality were recorded weekly for each group. Snails showing permanent reclusion into the inner shell whorl, absence of visible heartbeat for ten seconds, and/or shell discoloration were considered dead and removed without replacement. Only viable snails, showing heartbeats, actively crawling or attached to surfaces, or showing active body movement within the shell, were captured and measured. Growth was determined by measuring shell diameter. Reproductive output was determined by counting the number of egg masses, measuring egg mass length, and counting the number of embryos contained in each egg mass. Images were captured from snails and egg masses alongside a millimetre scale, using an Olympus SZX10 stereomicroscope equipped with an Olympus SC50 camera (Olympus, Japan), using the Olympus cellSens imaging software v2.3, with measurement via ImageJ software v1.54d [31]. Since snail mortality may have impacted total egg mass production per diet, we estimated weekly average egg mass production per snail. For each diet, snail survivors in a week with non-zero egg mass count were considered the egg-laying snails for that week, with the assumption that all snails for such week may have contributed to egg laying. In each diet, snail counts in weeks with no egg mass production were excluded.

All phenotypic data showing normal distribution and equal variance (Shapiro–Wilk’s *p* > 0.05 and Levene’s *p* > 0.05) were analysed using parametric one-way Analysis of Variance (ANOVA) test, followed by Tukey’s HSD test at 95% confidence level with Bonferroni’s correction for pairwise comparisons. When data were not normally distributed, non-parametric Kruskal–Wallis test followed by Dunn’s post-hoc test was adopted. Snail survival rates were compared using Log-rank (Mantel–Cox) test with Bonferroni’s correction.

### RNA isolation, library preparation, and RNA sequencing

Six size-matched (7–9 mm) snail samples from FF and SG groups, and all five survivors from the FL group were collected for transcriptomic analysis. Snails were gently rinsed in distilled water, and the shell surfaces were wiped with 70% alcohol before removal of snail tissue from the shell under sterile conditions. Individual snail tissues were placed into separate 2 mL tubes, immediately flash-frozen in liquid nitrogen and stored at –80 □ until further processing. To isolate total RNA, frozen snails were individually homogenised in 1 mL TRIzol reagent (Invitrogen) using an automated TissueLyser (Qiagen). Each homogenate was extracted with 200 µL chloroform as described by the manufacturer’s protocol with the resulting pellet dissolved in 40 µL RNase-free water. Residual genomic DNA was removed using a DNA-free kit (Invitrogen). Before and after DNAse treatment, the samples were examined for both quality and quantity using a spectrophotometer (DeNovix DS-11 FX). All RNA samples yielded a purity value (260/230 and 260/280 absorbance ratios) of ~ 2.0 and were diluted to 100 ng/µl in 10 µL of RNAse-free water. Illumina library preparation and sequencing was carried out at the Queen’s University Belfast Genomics and Cytometry Core Technology Unit (GCCTU). For quality control, an Agilent 4200 TapeStation Analyser v5.1 (Agilent Technologies) was used to assess the RNA integrity, and gDNA contamination check was performed using Agilent 5200 Fragment Analyser v3.1.0.12 (Agilent Technologies). RNA samples showed integrity (RIN) scores ranging from 8.1 to 10. Strand-specific mRNA-Seq libraries were prepared from the 17 samples using the KAPA mRNA HyperPrep Kit with PolyA enrichment for Illumina. Paired-end sequencing with 2×150 bp read length was performed on the 17 reverse-transcribed cDNA libraries using the Illumina NovaSeq 6000 platform.

### Bioinformatic analysis of Illumina RNA-Seq data

Quality control of raw FASTQ files used fastQC v0.11.8 (https://www.bioinformatics.babraham.ac.uk/projects/fastqc/), followed by multiQC v1.15 [32]. Raw FASTQ reads were trimmed for adapters using TrimGalore v0.6.10, yielding reads longer than 50 bp and with a Phred quality ≥ 30. *Biomphalaria glabrata* reference genome (xgBioGlab47.1) [33] and the associated GTF annotation (GCF_947242115.1) were downloaded from NCBI (https://ftp.ncbi.nlm.nih.gov/genomes/). Adapter-trimmed, high-quality reads were mapped to the *B. glabrata* reference genome using HiSAT2 v2.1.0 with default settings [34]. Samtools v1.9 [35] was used to convert SAM files to sorted BAM files. New GTFs for all samples were merged into a single matrix, and reads that uniquely mapped to a gene were counted using StringTie v1.3.6 [36].

### Differential gene expression analysis

Differential expression analysis was performed with DESeq2 package v1.42.1 [37] after initial removal of low expression genes with < 10 reads in more than five samples across the datasets. Inter-sample and inter-group variations were assessed via a sample-to-sample distance matrix and a principal component analysis (PCA), respectively, using the Euclidean algorithm [38]. Significance in the inter-group variation was tested using one-way Permutational Multivariate Analysis of Variance (PERMANOVA), which was performed using *adonis2* function of the vegan v2.7 package. Prior to distance matrix and PCA analyses, DESeq2 data was transformed to stabilise variance across the mean. For differentially expressed genes (DEGs), a subset of the DESeq2 result with adjusted *p*-values □ 0.01 and log_2_ fold change ≥ 1.0 or ≤ –1.0 was considered significantly differentially expressed. A global heatmap for DEGs across all snail samples was generated from normalised DESeq2 counts subjected to Z-statistics [37]. Differential gene expressions between the dietary treatment groups were examined in three pairwise comparisons: FL vs FF, FL vs SG, and SG vs FF. Estimates of DE counts that were unique to each diet group and those that overlapped between groups were computed. Heatmaps for pairwise comparisons of DEGs were generated with pheatmap package v1.0.13 using the top 50 hits, after the normalised count data were log-transformed and scaled.

### Identification and annotation of DEGs

Unique DEG IDs were mapped to the DAVID bioinformatics database [39]. For additional annotation, gene IDs were also mapped to the UniProtKB database [40]. Genes among the top-ranked DEGs and other genes of interest that were labelled as uncharacterised were further identified by (i) functional domain analysis using InterPro v102.0 with preconfigured cut-off thresholds [41], (ii) the *hmmscan* module of HMMER v3.4 [42] with the default gathering threshold, and (iii) by manual interrogation of their protein-coding nature on NCBI database (https://www.ncbi.nlm.nih.gov/).

### Functional enrichment analyses

The biological processes (BP), cellular components (CC), molecular functions (MF), and pathways associated with DEGs were identified using ShinyGO v0.80 [43]. This involved the mapping of all DEGs in each diet group to the AmiGO2 database [44] and Kyoto Encyclopaedia of Genes and Genomes (KEGG) database [45]. The filtered DESeq2 result containing genes with detectable expression (without the lowly-expressed genes) was used as background gene list. Enrichment *p*-values were determined using a hypergeometric test [46] and False Discovery Rate (FDR) values were calculated following the Benjamini-Hochberg method [47]. All functional enrichment analyses were performed at FDR ≤ 5%. To prevent enrichment masking, redundant gene ontology (GO) terms were programmatically removed. Further analysis was performed to identify gene sets that were upregulated and downregulated in the enriched GO terms and KEGG pathways.

### Microbial metatranscriptomic analysis

To investigate the effects of diet on the snail microbial community, we used trimmed FASTQ sequences that did not map to the *B. glabrata* genome to query microbial genomes based on *k*-mer matching using Kraken 2 database v2.1.3 [48]. We used a pre-built Kraken 2 database (PlusPF v2024) that contained viral, archaeal, bacterial, protozoan, fungal and human genomes. Kraken 2 analysis was followed by Bayesian re-estimation of species abundance, annotated to the class level, using Bracken v3.1 [49]. Both Kraken and Bracken analyses were performed on Galaxy (https://usegalaxy.eu) [50]. Bracken results were denoised by removing non-microbial taxa. Alpha diversity was analysed to quantify within-sample taxonomic diversity, with Observed and Chao1 metrics used to estimate taxonomic richness, Simpson index to determine the level of evenness or dominance in taxonomic distribution, and Shannon index for integrated information on taxonomic richness and distribution. To assess differential microbial composition between diet groups, beta diversity analysis was performed using Bray–Curtis dissimilarity index and one-way PERMANOVA. To validate PERMANOVA assumptions, a follow-up beta-dispersion test was executed.

Relative abundance (RA) of microbial taxa across the groups was estimated using the total sum scale-transformed OTU data treated with Kruskal–Wallis method with FDR correction. Only taxa with significant RA (FDR ≤ 0.05) were filtered and pairwise Mann– Whitney *U* (Wilcoxon rank-sum) test was performed to compare intergroup differences in significance. Since the RA approach is compositional and bias-prone, we further performed differential abundance (DA) analysis using the ANCOM-BC v2.6.0 pipeline which addresses compositionality and bias [51]. Based on rare taxa filtering (taxa with < 3 non-zero samples) and microbiome library size, parameters including *prv_cut* = 0.17647 and *lib_cut* = 100,000 were used for ANCOM-BC analysis, with FL as baseline. Furthermore, we statistically determined taxa that were strongly associated with, but not necessarily abundant in, each diet group, using *multipatt* function of the Indicspecies v1.8.0 package. Indicator cut-off value (IndVal index) was set at ≥ 70, and significant associations were reported at *p* < 0.01.

### Correlation analysis between snail microbiome and DEGs

We further asked if dietary regulation of gene expression correlated with microbial enrichment. To reduce noise and dimensionality, we focussed majorly on the relatively or differentially significant taxa. Total-sum scaled OTU matrix was transformed by centred log-ratio (CLR) with zero-replacement using multiplicative method, followed by Mantel statistics and Spearman correlation with 999 permutations. Through monotonic taxon–gene analyses, we also used subsets of DEGs associated with growth, immunity and reproduction to explore potential correlation between microbial taxa and the snail physiology.

### Software analysis and visualisation

Phenotypic, transcriptomic, and metatranscrcriptomic data were analysed and visualised using R statistical packages (R v4.3.2) [52]. The KEGG and GO enrichment pathways, and chord plots were visualised using SRplot [53]. The general methodological pipeline of the study is summarised in Fig. 1.

**Fig. 1.**
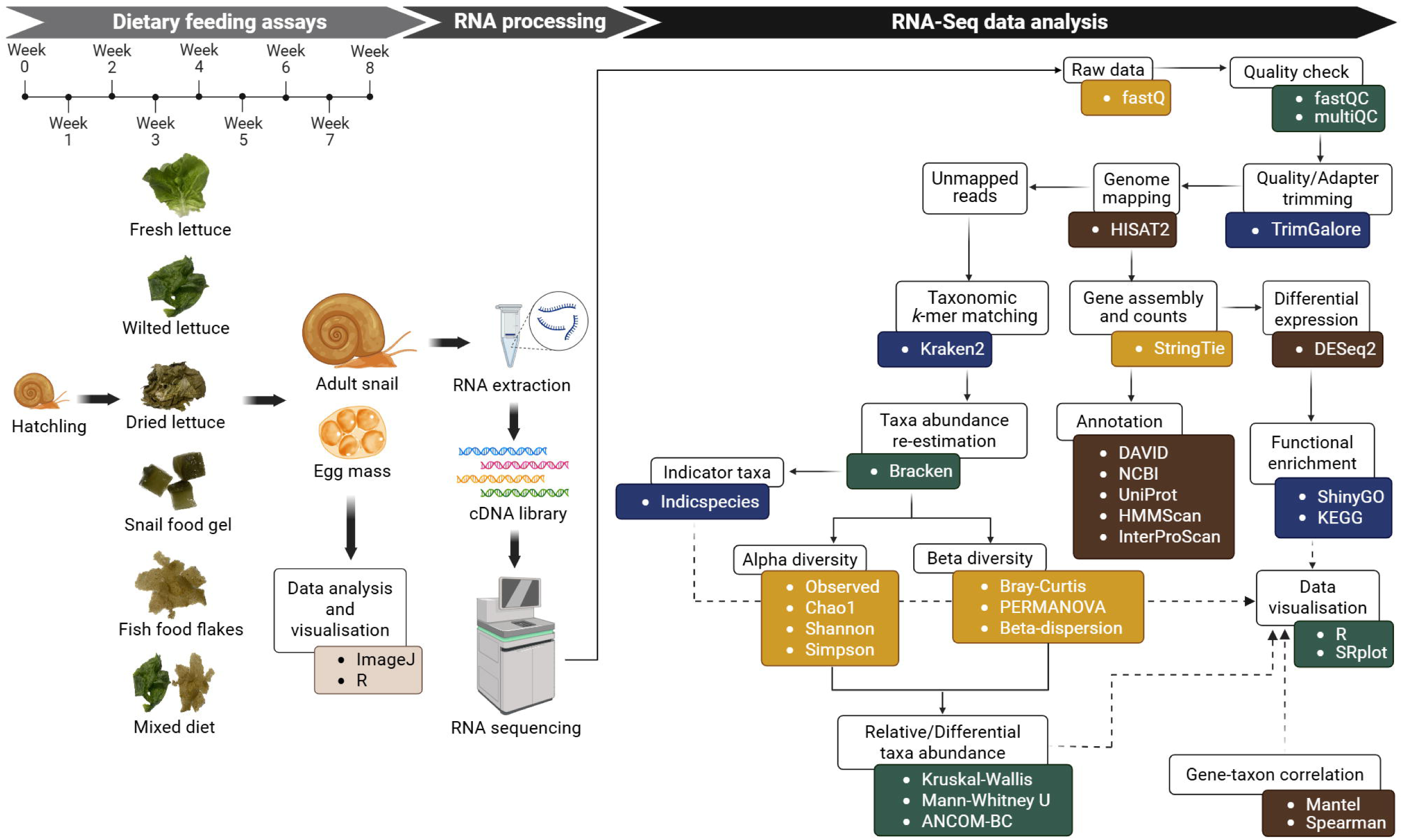
Methodological flow chart. The effects of diets on snail survival, growth and reproductive phenotypes were examined, with 8-week-old adult snails selected for RNA sequencing for transcriptomic and metatranscriptomic analyses.

## Results

### Diet influences key phenotypes in *B. glabrata*

As shown in Table 1 and Fig. 2, FF and SG diets promoted increased growth, survival and fecundity compared to lettuce diets that showed the slowest growth and lowest survival rates. After eight weeks of feeding, FF and SG snails had grown significantly larger than FL snails (*p* < 0.001) (Table 1 and Fig. 2A). Dried lettuce (DL) also supported growth more effectively than FL (*p* = 1.2e-04) (Table 1 and Fig. 2A).

**Fig. 2.**
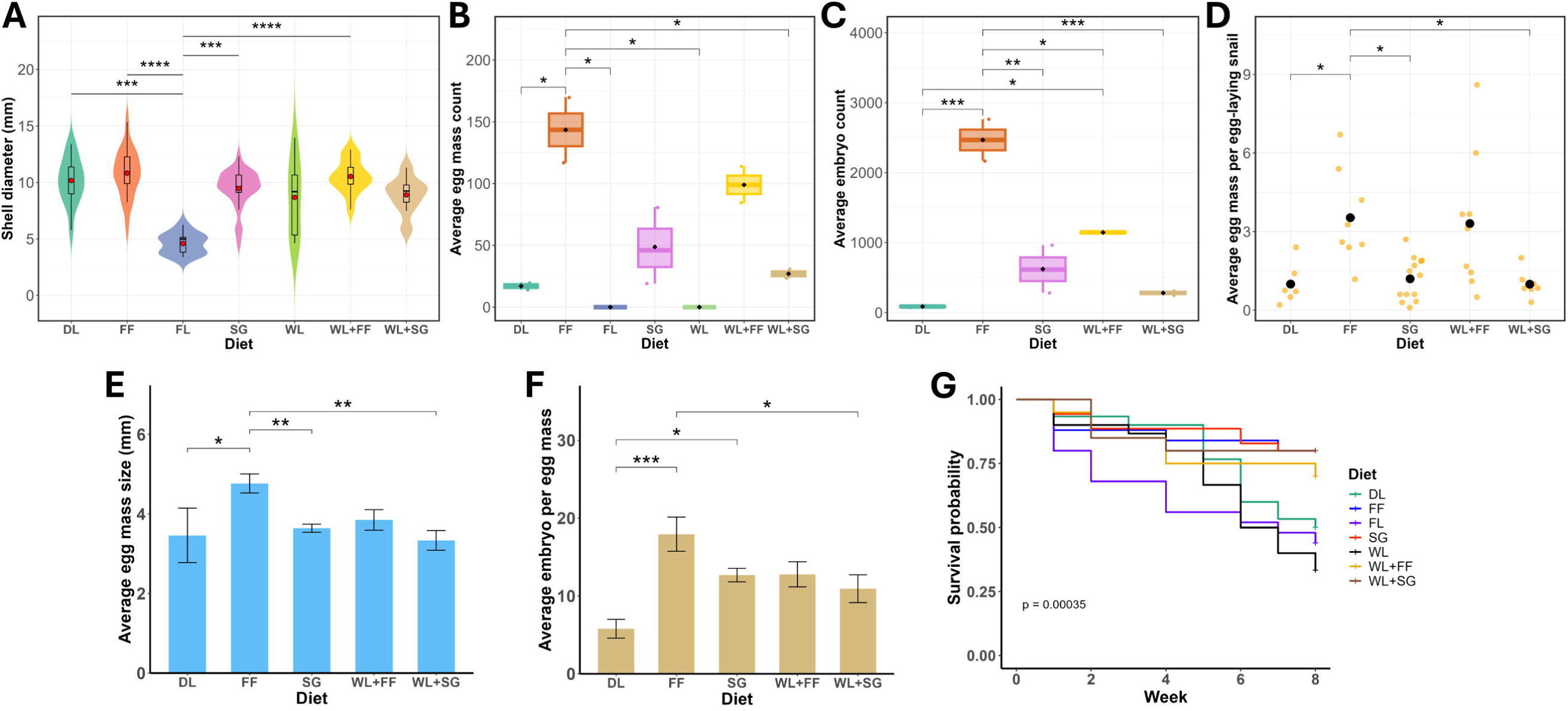
Nutrition impacts growth, reproduction and survival phenotypes in *Biomphalaria glabrata*. (A) Growth, based on shell size as measured after 8 weeks of feeding. (B) Reproduction, based on number of egg masses produced per diet within 8 weeks. (C) Reproduction, based on number of embryos produced per diet within 8 weeks. (D) Reproduction, based on average number of egg masses produced per egg-laying snail per diet within 8 weeks. Each yellow jittered point represents an average number of egg mass produced per snail in a particular week. (E) Reproduction, based on average size of egg masses produced per diet within 8 weeks. (F) Reproduction, based on average number of embryos per egg masses per diet within 8 weeks. (G) Survival, Kaplan–Meier survival curve showing percent survival for an 8-week period. In Figs. A–D, central red or black dots represent mean value of the data. The error bars in Figs. B–F represent standard error of the mean (SEM). DL, dried lettuce; FF, fish food flakes; FL, fresh lettuce; SG, snail gel. In all relevant Figures, \**p*adj ≤ 0.05, \*\**p*adj ≤ 0.01, \*\*\**p*adj ≤ 0.001, \*\*\*\**p*adj ≤ 0.0001.

**Table 1.**
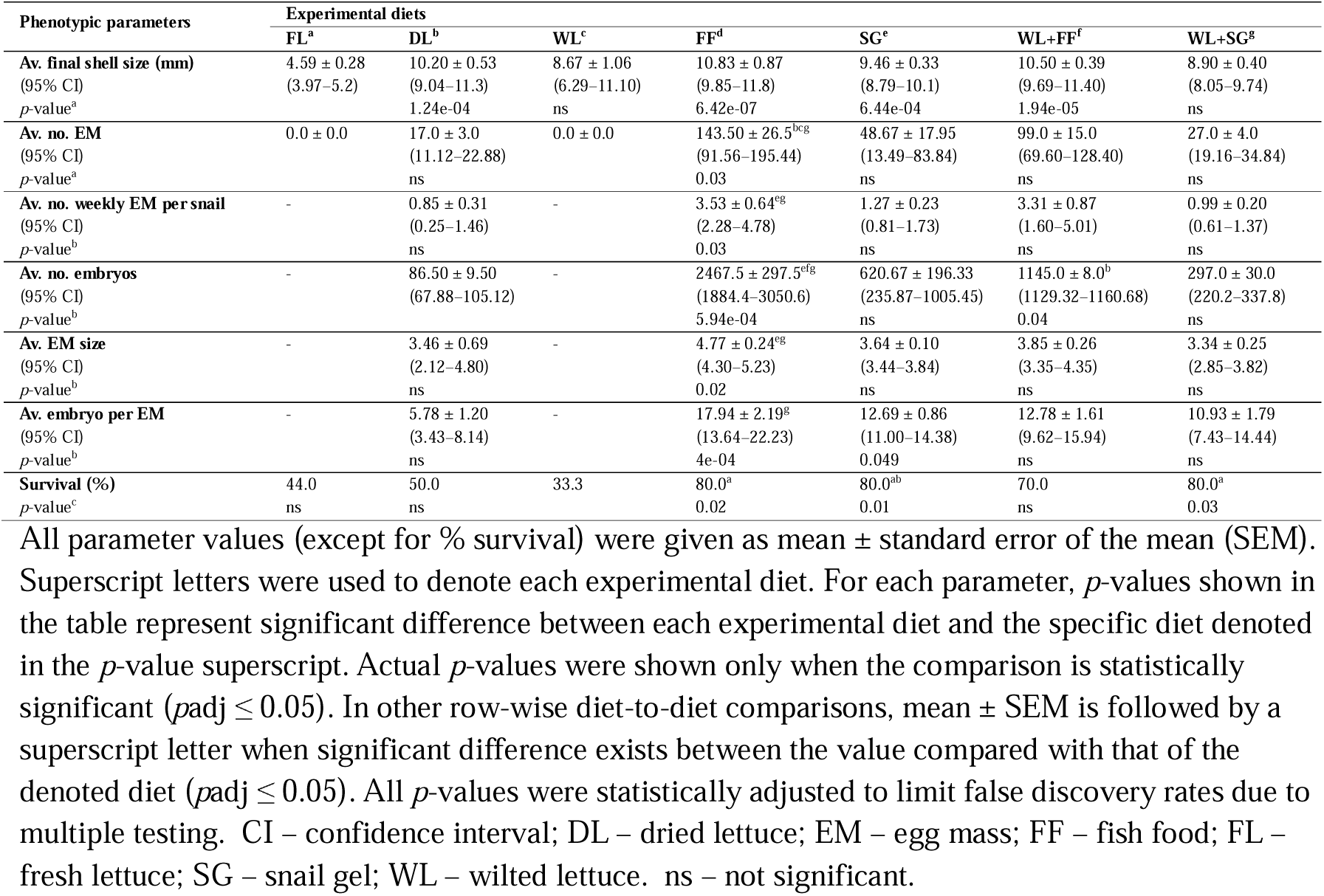
Statistical data on snail phenotypes after 8 weeks of experimental feeding.

In measurements of reproduction, neither FL nor WL produced any egg masses within the feeding period (Table 1 and Fig. 2B), while FF and SG supported earlier onset of sexual maturity, commencing egg-laying within four to five weeks of hatching. Snails fed on FF produced more egg masses and embryos than any other group (Table 1, Figs. 2B and C). In contrast to FL and WL, DL did support egg mass production, although the difference was not statistically significant (*p* > 0.05) (Table 1, Figs. 2B and C). FF-fed snails also laid significantly larger egg masses than other diet groups, except for dual WL+FF-fed snails (Table 1 and Fig. 2E). Both in terms of total embryo output and average embryo count per egg mass, FF produced more embryos than other diets (Table 1). The average embryo count per egg mass in FF was more significant than in DL (*p* = 4e-04) and WL+SG (*p* = 0.04). Also, SG produced significantly higher number of embryos per egg mass than DL (*p* = 0.049) (Table 1 and Fig. 2F).

Fish food (FF), SG and WL+SG similarly supported higher snail survival (80% at eight weeks) than the lettuce diets where survival rates ranged between 33.3% and 50%. Wilted lettuce (WL) gave the lowest chances for survival (Table 1 and Fig. 2G). Log-rank test showed a statistically significant difference in survival curves across all diet groups (*p* = 0.00035) (Fig. 2G). Pairwise Log-rank test also showed significant differences in survival rates between diet groups (Table 1). Weekly mortality rate per diet did not appear to skew egg mass production. Although egg mass production per snail generally increased with time from the onset of sexual maturity, each FF snail laid an average of 3.5 egg masses per week, followed by WL+FF (3.3), SG (1.3), WL+SG (1.0), and DL (0.85) (Fig. 2D). The average egg mass output per egg-laying snail was significantly different among diet groups (Kruskal– Wallis: *stat* = 16.8, *df* = 4, *p* = 0.002) (Table 1 and Fig. 2D), but still in a similar pattern to the mortality-unadjusted egg-laying data (Fig. 2B). Supplementing WL with either FF or SG showed clear improvements in all phenotypic parameters tested. For example, WL+FF and WL+SG diets increased survival, by 36.7% and 46.7% respectively, and egg mass production when compared with WL alone (Table 1).

### Transcriptome data overview

Sequenced cDNA libraries each returned between 43.0–65.5 million reads having between 5.6–8.7 billion base pairs (Additional file 3: Table S1). After stringent quality and adapter trimming, all (100%) reads and an average of 97.8% of the total base pairs remained, with average read length of 130 bp. Libraries contained between 41.6–64.1 million unique mappable reads. Read quality and mapping data per sample are presented in Additional file 3: Table S1. Average metrics generated per diet group are presented in Table 2.

**Table 2.** Average RNA-seq data quality and mapping metrics per diet group.

|  | FF | FL | SG |
| --- | --- | --- | --- |
| Libraries (*n*) | 6 | 5 | 6 |
| Total reads (pre- and post-trimming) | 59,387,162 | 49,538,561 | 55,681,557 |
| Reads with adapters (%) | 23.44 | 25.02 | 23.94 |
| Base pairs processed | 7,847,318,010 | 6,492,614,005 | 7,298,807,227 |
| Base pair retained post-trimming (%) | 98.18 | 97.25 | 97.80 |
| GC content (%) | 38.66 | 36.33 | 38.83 |
| Read length (bp) | 131 | 129 | 130 |
| Mappable reads | 58,077,978 | 47,885,890 | 54,188,096 |
| Reads mapped to the reference genome (%) | 82.92 | 67.33 | 79.93 |
| Reads unmapped to the reference genome (%) | 17.08 | 32.67 | 20.07 |
| Unmapped reads classified by taxonomy (%) | 14.83 | 18.59 | 16.73 |

“bp” = base pairs. Bp processed indicates the complete length of sequence processed by TrimGalore. Read length is the average length of each read after trimming, as estimated by multiQC.

### Diet influences differential expression of genes

Transcriptome assembly (Additional file 4: Dataset 1) followed by DESeq2 filtering identified a total of 21,887 expressed genes across all libraries (Additional file 5: Dataset 2). PCA analysis showed a significant inter-group variation in gene expression patterns (PERMANOVA: *R^2^* = 0.48, *F* = 6.49, *p* = 0.001) (Fig. 3A). However, SG and FF snails were more similar in their gene expression patterns than the more distant FL group (Figs. 3A and B; Additional file 1: Fig. S1). In the FL vs FF comparison, 4,114 genes were differentially expressed (Figs. 3C and D; Additional file 6: Dataset 3). FF-upregulated DEGs (*p* < 0.01) included bactericidal permeability-increasing protein (BPI, also known as lipopolysaccharide-binding protein, LBP), mucin-5AC (MUC5AC), temptin (TEMPT), aplysianin-A (APLY-A), hemocyanin 1 (HCY1), serine protease inhibitors (e.g. SERPINB3, SERPINB6, SPN42Dd), adhesive plaque matrix protein (FP1), and lectins (e.g. CLEC) (Fig. 3G; Additional file 7: Table S2). Meanwhile, FF-downregulated DEGs (*p* < 0.01) included glutathione transferases (GST3, GST7), 1-deoxyxylulose-5-phosphate synthase YajO, pirin (PIR), C-factor, oxidoreductase HTATIP2, and FMN-dependent NADPH-azoreductase (AZR) (Fig. 3G; Additional file 7: Table S2).

**Fig. 3.**
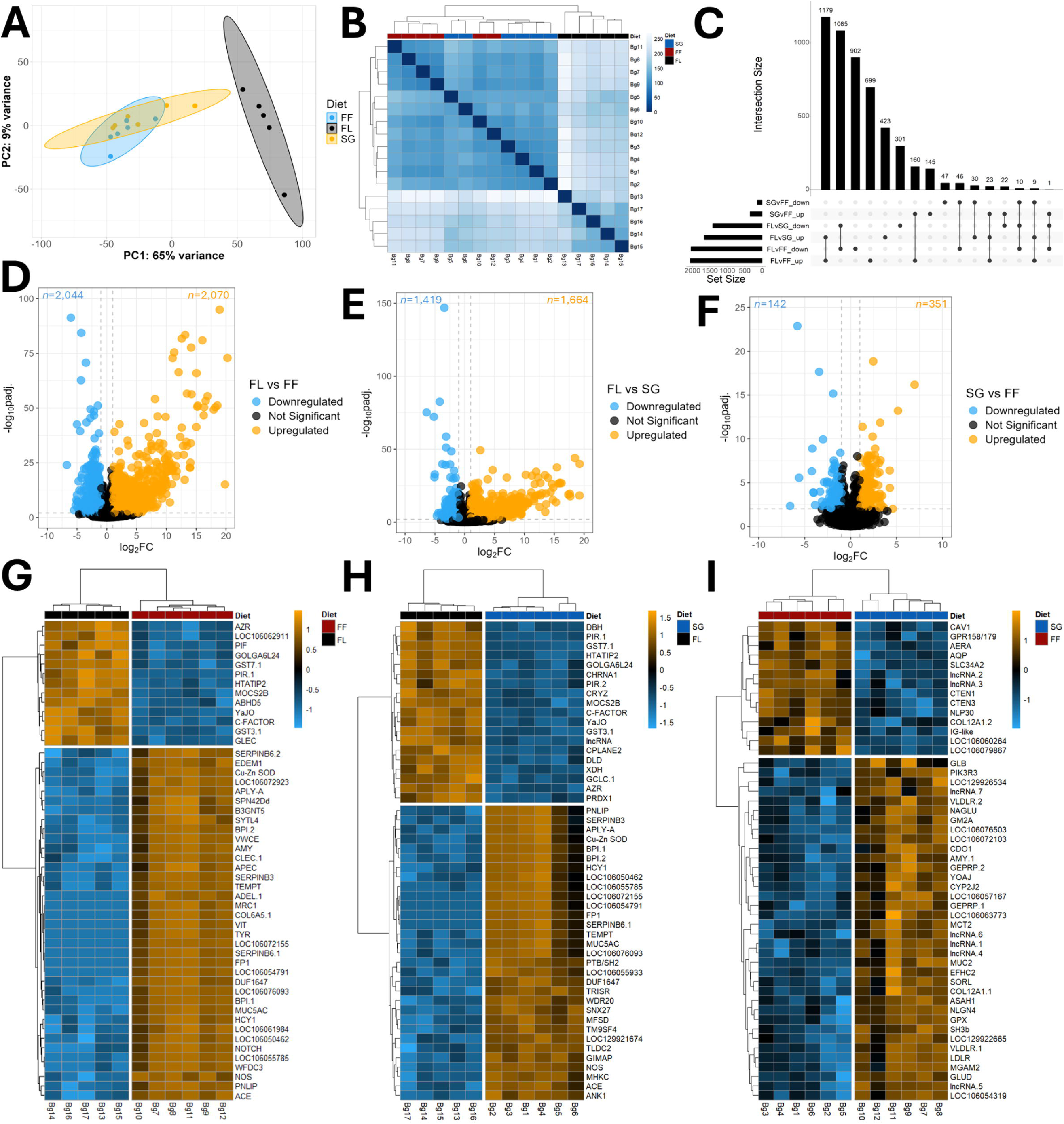
*Biomphalaria glabrata* gene expression patterns are impacted by nutritional intake, with largest gene expression differences in comparisons between fresh lettuce (FL), and fish food (FF) or snail gel (SG). (A) Principal component analysis (PCA) and (B) Sample-to-sample clustering using Euclidean distance matrix (Darker blues indicate higher correlation in gene expression pattern between two snail samples on the x and y axes) both highlight diet-specific patterns of differential gene expression; (C) Upset plot of differentially expressed genes (DEGs, *p* < 0.01) across all dietary comparisons indicates that the highest number of DEGs are found in comparisons between FL and FF or SG. Horizontal bars (set size) display the total number of DEGs in each comparison. Vertical bars (intersection size) indicate the number of DEGs that are uniquely expressed or shared among diet groups. The matrix below the vertical bar plots indicates the set of genes represented in each bar. (D– F) Volcano plots of significant differentially expressed genes in FL vs FF, FL vs SG, and SG vs FF comparisons, respectively. Plots highlight DEGs within the ranges –log_10_*p*adj > 2 and log2 fold change (log_2_fc) > 1 or < –1. The x-axis (log_2_fc) presents the expression level and the y-axis (– log_10_*p*adj) presents the statistical significance of the differential expression. Each point represents a unique gene. (G–I) Pairwise heatmap visualisation of top 50 most significant differentially expressed genes (ranked by adjusted p-value) in FL vs FF (G), FL vs SG (H), and SG vs FF (I) comparisons. Each row represents a gene, each column represents a snail sample, and each cell represents the scaled transformed normalised value for gene expression. Lighter brown represents higher upregulation while lighter blue colour represents higher downregulation. Genes are clustered hierarchically based on average agglomeration and Euclidean distance measure. The hierarchical dendrogram indicates similarities among samples based on the normalised gene expression values. Bg1–Bg6 were SG snails; Bg7–Bg12 were FF snails; Bg13–Bg17 were FL snails. Full gene names and gene IDs of all DEGs are documented in Additional file 6: Dataset 3.

In the FL vs SG comparison, 1,664 (54%) of the 3,083 significant DEGs (*p* < 0.01) were upregulated, while 1,419 (46 %) genes were downregulated (Figs. 3C and E, Additional file 6: Dataset 3). As expected, SG and FF shared more similar sets of DEGs (Figs. 3C and H, Additional file 7: Table S2), with just 493 genes DE in the SG vs FF comparison. Of these, 351 (71%) genes were upregulated and 142 (29%) were downregulated in the FF snails (Figs. 3C and F, Additional file 6: Dataset 3). Among the FF upregulated genes were glutamate dehydrogenase (GLUD), lipoprotein receptors (LDLR and VLDLR), globin-repeat domain-containing protein or globin-like oxygen transporter (GLB), alpha-amylase (AMY), cysteine dioxygenase type 1 (CDO1), and EF hand 2 domain-containing protein (EFHC2). By contrast, genes encoding aquaporin (AQP), ctenidin (CTEN), aerolysin (AERA), sodium-dependent phosphate transport protein 2B (SLC34A2) among others, were downregulated in SG vs FF comparisons (Fig. 3I). Among the top DEGs were several lncRNAs (Additional file 6: Dataset 3; Additional file 7: Table S2).

### Diet enriches specific metabolic pathways and gene sets

Gene ontology (GO) annotations showed that metabolic processes involving fatty acid and lipid oxidation were more enriched in FF snails relative to FL, while glycoprotein biosynthesis and metabolism were more prominent in SG (Figs. 4A–C). The FF vs SG comparisons highlighted several shared signals, including dynein light intermediate chain binding, minus-end-directed microtubule motor activity, and microtubule motor activity (Figs. 4A and B). Conversely, mechanisms and cellular components uniquely involved in FL-fed snails differed widely. These include glutathione metabolism, caveola assembly, (nc)RNA processing, and activities involving cytokines, hormones, signalling receptors, and receptor ligands (Fig. 4C; Additional file 8: Dataset 4). KEGG pathway analysis (Fig. 4D) highlighted that fatty acid degradation, glycosaminoglycan degradation, butanoate (butyrate) metabolism and propanoate (propionate) metabolism among others, were associated with FF. Snails maintained on SG showed KEGG links to biosynthesis of various glycans, especially mucin type O-glycan and type N-glycan, while FL snails were dominated by KEGG mechanisms linked to metabolism of xenobiotics, glutathione, drug, and neuroactive ligand-receptor interactions (Fig. 4D; Additional file 9: Dataset 5).

**Fig. 4.**
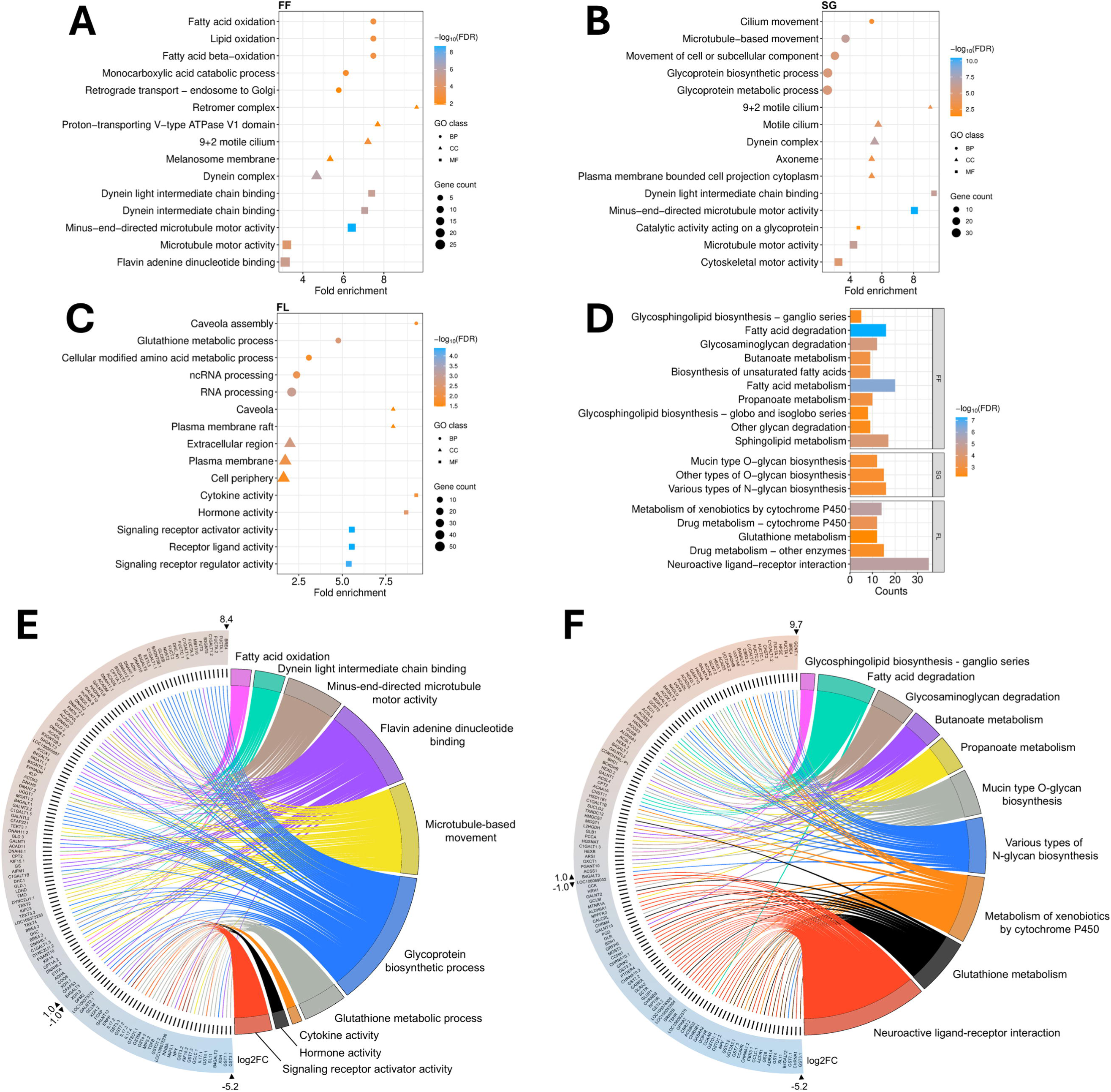
GO and KEGG enrichment analysis highlights diet-related overrepresentation in GO terms and KEGG pathways and differentially expressed gene sets connected with each term and pathway in *Biomphalaria glabrata*. (A–C) Top 5 GO terms for fish food (FF; A), snail gel (SG; B) and FL (FL; C)-fed snails, within biological process (BP), cellular component (CC) and molecular function (MF) classes. (D) Top 10 KEGG pathways for FL, FF and SG groups. For A–D, overrepresented GO terms and KEGG pathways were first filtered to remove terms with false discovery rate (FDR) higher than 0.05; and then sorted by fold enrichment value from highest to lowest in each diet group. Only KEGG pathways with FDR ≤ 0.05 in each treatment were shown in (D). The fold enrichment value indicates the ratio of genes overrepresented in a specific pathway to the ratio of the same pathway genes in the background gene list. Higher FDR values suggest higher likelihood of the pathway occurring by chance under the specific dietary treatment; (E) Selected top and functionally relevant GO terms and (F) Selected top and functionally relevant KEGG pathways, in fish food (FF) and or snail gel (SG) groups, in comparison with FL. On the gene list arch, log_2_fc < –1.0 indicates downregulation and > 1.0 indicates upregulation in FL comparisons with either FF or SG. All genes listed were differentially expressed with *p* < 0.01. Full gene names and gene IDs of all DEGs are documented in S4 Table.

Differentially expressed gene sets in some selected GO and KEGG enrichment indicators are presented in Figs. 4E and F. In the FF group, for instance, transcripts encoding trifunctional enzyme subunits (HADHA and HADHB), mitochondrial 3-ketoacyl-CoA thiolase (ACAA2), carnitine O-palmitoyltransferases (CPTs), acyl-CoA dehydrogenases (ACADL, ACADVL), peroxisomal acyl-coenzyme A oxidases (ACOX1 and ACOX3), and another peroxisomal bifunctional enzyme (EHHADH) were among the upregulated gene sets that may have significantly facilitated fatty acid oxidation/degradation (Figs. 4E and F). Our data captured ten genes in the butanoate metabolism pathways (HADHA, HADH, EHHADH, ACADS, L2HGDH, BHD1, ALDH5A1, OXCT1, HMGCS1, BDH1) and eleven genes in the propanoate pathway (ACOX1, ACOX3, HADHA, EHHADH, ACADS, ALDH6A1, ACSS1, ACSS3, SUCLG2, PCCA, BCKDHB) (Fig. 4F).

The observed KEGG glycoprotein biosynthesis pathways (including mucin type O-glycan and N-glycans; Fig. 4D) in SG snails reflects transcriptional upregulation of beta-1,4-N-acetylgalactosaminyltransferase (BRE4), some members of beta-1,3-galactosyl-O-glycosyl-glycoprotein beta-1,6-N-acetylglucosaminyltransferases (e.g. GCNT.1, GCNT.2, and GCNT2), glycoprotein 3-alpha-L-fucosyltransferase A-like genes (e.g. FUCTA.1, FUCTA.2), glycoprotein-N-acetylgalactosamine 3-beta-galactosyltransferase 1-like genes (e.g. C1GALT1.1, C1GALT1.2, C1GALT1.3), and beta-1,3-galactosyltransferases (e.g. B3GALT5 and B3GALT2). Genes including B4GALT2 and GALNT13 were downregulated in this process (Fig. 4F). Signals that were enriched in FL snails were downregulated in FF and SG. Metabolism of xenobiotics by cytochrome P450 and glutathione metabolism were among the prominent signals in the FL snails (Fig. 4D), with glutathione transferases (GST3.1, GST3.2, GST4, GST6, GST7.1 and GST7.3) including glutathione transferase omega-1 (GSTO1.1 and GSTO1.2) amongst the most highly regulated genes (Figs. 4E and F).

### Metatranscriptomics suggests that diet alters the structure of the *B. glabrata* microbiome

The microbiome library size per sample ranged between 138,472 and 314,209 sequence reads mapping to microbial genomes (Additional file 2: Fig. S2). A total of 104 classes of microbes (2.9% yeasts, 8.7% viruses, 8.7% protists, 11.5% archaea, and 68.3% bacteria) were identified transcriptionally across the samples (Fig. 5A; Additional file 10: Dataset 6), with *Gammaproteobacteria* being the most predominant taxon (Fig. 5D). While Simpson diversity showed significant difference in taxonomic class-level distribution between FF and SG groups (*p* = 0.047), other alpha diversity metrics (Observed Richness, Chao1, and Shannon) were not significantly different across the groups (*p* > 0.05) (Fig. 5B). Beta diversity analysis using Bray–Curtis dissimilarity (Fig. 5C) revealed significant differences in microbial composition across the diet groups (PERMANOVA: *R^2^* = 0.37, *F* = 4.17, *p* = 0.001). Pairwise PERMANOVA comparisons between groups further confirmed significant inter-group differences (FL vs FF: *R^2^* = 0.28, *F* = 3.54, *p*adj = 0.01; FL vs SG: *R^2^* = 0.29, *F* = 3.78, *p*adj = 0.01; SG vs FF: *R^2^* = 0.41, *F* = 6.94, *p*adj = 0.006). The higher F-values suggested that the observed inter-group differences in microbial community were substantially larger than the sample-to-sample variation within each group. However, significant beta-dispersion (*F* = 4.63, *p* = 0.03) indicated that the observed PERMANOVA results may be partly influenced by within-group variations (Fig. 5C). Taken together, these results suggested that dietary changes induced compositional shifts in microbiome between groups (beta diversity) while the internal taxonomic richness in each sample did not vary significantly across the groups (alpha diversity).

**Fig. 5.**
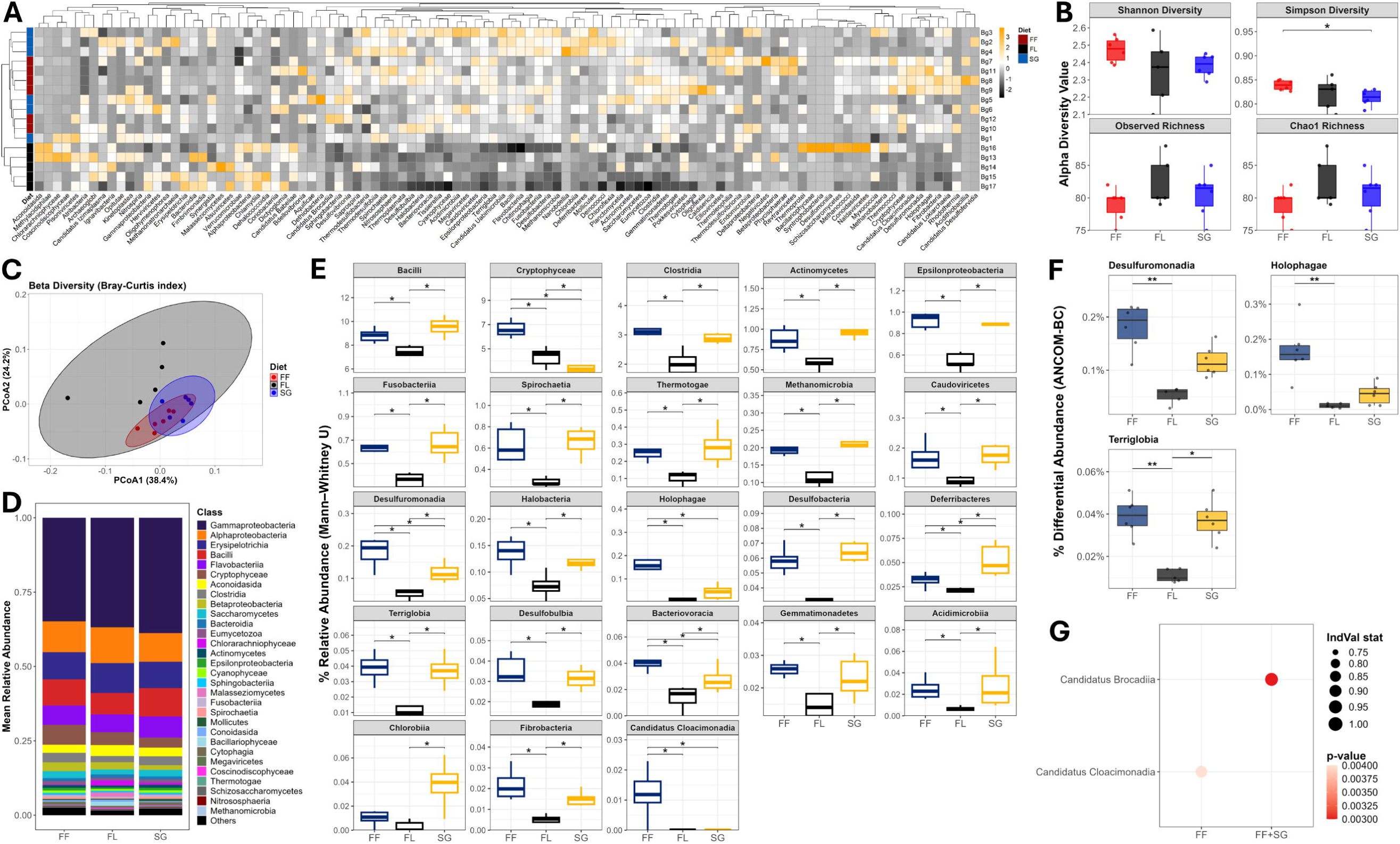
Diet induces changes in *Biomphalaria glabrata* microbiome diversity. (A) Heatmap showing the relative abundance in microbiome OTUs (operational taxonomic units) across samples. By our study approach, OTUs were taxonomically assigned classes with abundance counts derived from Bracken classification. OTUs were normalised by total sum scaling. The colour scale represents Z-scores across samples. At Z = 0, the sample has mean-level abundance for that OTU. Z > 0 means above-average abundance for that OTU, while Z < 0 means below-average abundance for that OTU. Snail gel (SG)-fed snails (Bg1–Bg6); fish food (FF)-fed snails (Bg7–Bg12); fresh lettuce (FL)-fed snails (Bg13–Bg17). (B) Alpha diversity metrics showing Shannon index, Simpson index, Observed taxonomic richness, and Chao1 richness in microbial diversity across samples. (C) Principal coordinate analysis (PCoA) representation of Bray–Curtis distance showing beta diversity in microbiome structures of the three diet groups. (D) Stacked bar plots showing top 30 most dominant taxa across diet groups. Class legend was sorted in descending order of dominance. (E) Comparisons of significant relatively abundant taxa between diet groups, \**p*adj < 0.05. (F) ANCOM-BC comparisons of significant differentially abundant taxa between groups, \**p*adj < 0.05, \*\**p*adj < 0.01. (G) Bubble plot showing multilevel pattern analysis for indicator taxa across diet groups. IndVal stat represents the mean value of specificity (taxon occurs mostly in that group) and fidelity (taxon occurs in most samples of that group). IndVal stat ≥ 0.7 suggests a stronger indicator taxon and *p* < 0.01 indicates higher significance of the diet–taxon association.

Our pairwise comparisons showed that 23 microbial classes were relatively more abundant between diet groups (Fig. 5E; Additional file 11: Table S3). *Bacilli*, *Clostridia*, *Actinomycetes*, *Epsilonproteobacteria*, *Fusobacteriia*, *Spirochaetia*, *Thermotogae*, *Methanomicrobia*, *Caudoviricetes*, *Halobacteria*, *Desulfobacteria*, *Terriglobia*, *Desulfobulbia*, *Gemmatimonadetes*, *Acidimicrobiia* and *Fibrobacteria* were more relatively abundant in both FF and SG groups than the FL group, while *Cryptophyceae*, *Desulfuromonadia*, *Holophagae*, and *Bacteriovoracia* members were relatively more abundant in FF snails than other groups. Only *Deferribacteres* bacteria were more relatively enriched in SG than other groups, while members of *Cryptophyceae* were more highly represented in FL than SG snails (Fig. 5E). Ninety-five (95) of the total 104 taxa identified passed ANCOM-BC filtering (Additional file 12: Dataset 7). After compositional bias correction, three taxa (*Desulfuromonadia*, *Holophagae* and *Terriglobia*), which are also among the relatively abundant taxa, remained differentially abundant across diet groups (*q* < 0.05) (Fig. 5F; Additional file 12: Dataset 7). This suggested that at least three out of the relatively abundant taxa were truly differentially abundant between diet groups. Two indicator taxa in the *Candidatus* cluster were uniquely and consistently associated with FF and SG diets. *Candidatus Cloacimonadia* was strongly associated with FF (*stat* = 0.87, *p* = 0.004), while *Candidatus Brocadiia* (*stat* = 0.94, *p* = 0.003) showed strong association with both FF and SG snails (Fig. 5G; Additional file 13: Text S1).

### Diet-induced shift in microbiome structure strongly correlates with differential gene expression

A Mantel test between the OTU distance matrix of the significant microbial taxa and the DEG distance matrix in the FL vs FF comparison showed a significant relationship between these measures (*r* = 0.63, *p* = 0.001) (Fig. 6A). Strong Mantel associations were similarly observed between microbial taxa and DEG subsets involved in growth, immunity and reproduction (r ≥ 0.69, *p* = 0.001) (Fig. 6 B–D). In agreement with the ANCOM-BC model result, *Holophagae*, *Desulfuromonadia* and *Terriglobia* were among the top taxa having strong positive Spearman correlation with upregulated DEGs in FF-fed snails (Fig. 6 E–G). *Holophagae* bacteria were most positively correlated with growth and reproduction (Figs. 6E and G), and *Spirochaetia* correlated most positively with immunity (Fig. 6F). The two indicator taxa, *Candidatus Cloacimonadia* and *Candidatus Brocadiia* also showed strong and significant positive correlation with growth, immunity and reproduction in the FF-fed snails. Besides the bacterial taxa, members of the class *Caudoviricetes* (tailed phages) also showed positive associations with snail immunity and growth (Figs. 6E and F). In contrast to these observations, *Chlamydiia* showed inverse correlation with the upregulated DEG subsets in FF snails, especially in regards to immunity-related transcripts (Fig. 6F).

**Fig. 6.**
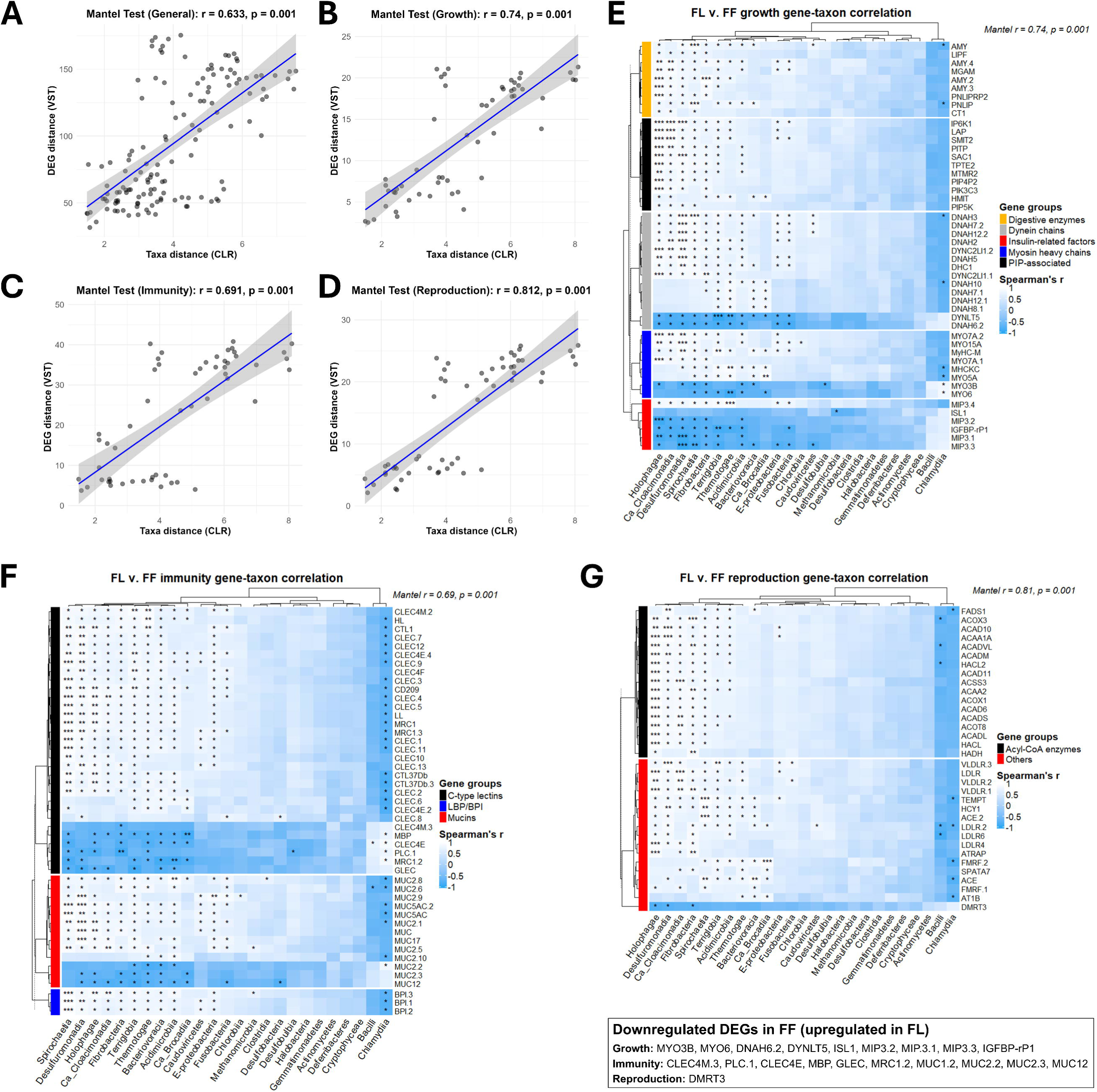
Diet-induced gene expression pattern significantly correlates with microbiome diversity in *Biomphalaria glabrata*. (A–D) Fresh lettuce (FL) vs fish food (FF) Mantel test scatter plots showing the relationships between (A) the variance-stabilised transformed (VST) values of all differentially expressed genes (DEGs) and centred log-ratio (CLR) values of significant microbial OTUs. (B) Significant taxa and growth-related DEG set. (C) Significant taxa and immunity-related DEG set. (D) Significant taxa and reproduction-related DEG set. In A–D, each geometric black point indicates DEG-taxon correlation. (E–G) FL vs FF heatmaps illustrating the Spearman correlation between (E) Significant taxa and growth-related DEG set. Heatmaps were generated using ComplexHeatmap package v2.18 (F) Significant taxa and immunity-related DEG set. (G) Significant taxa and reproduction-related DEG set. *Ca_Brocadiia – Candidatus Brocadiia*; *Ca_Cloacimonadia – Candidatus Cloacimonadia*; *E-proteobacteria – Epsilonproteobacteria*; PIP – Phosphatidylinositol phosphate; LBP/BPI – Lipopolysaccharide-binding protein/Bactericidal permeability-increasing protein; Acyl-CoA – Acyl coenzyme A. On Spearman’s r scale legend, lighter shades (> 0) indicate positive or direct correlation while darker shades (< 0) indicate negative or inverse correlation. Each cell on the heatmap suggests the degree of correlation between the two variables on the row and column, with \**p*adj < 0.05, \*\**p*adj < 0.01, \*\*\**p*adj < 0.001. Full gene names and gene IDs of all DEGs are documented in S4 Table.

## Discussion

While previous studies highlighted the physiological impacts of various dietary sources on *Biomphalaria* snails, this study is the first to examine the molecular mechanisms of diet-induced phenotypes. Our study has employed RNA-Seq methods to profile the impacts of dietary variation on the *B. glabrata* transcriptome and metatranscriptome, in relation to snail survival, growth and reproduction phenotypes.

We showed that feeding *B. glabrata* on an exclusive FL diet does not optimally support survival, growth or reproduction, despite lettuce being a widely used diet for laboratory snail maintenance. Our observations on diet/phenotype associations are consistent with some previous reports [12, 20]. Caloric contents of FF, FL and SG food sources correlated with the observed phenotypes; Lettuce represents a low-calorie diet, supplying 0.16 kilocalories per gram (kcal/g) (producer’s nutritional information), while the FF diet provided 3.96 kcal/g (manufacturer’s nutritional information). Our prepared SG diets was also calorific as it contained FF flakes, wheatgerm (3.85 kcal/g), barley grass (3.10 kcal/g) and whole milk powder (5.15 kcal/g). Higher fecundity performance of FF diet may be attributed to its fat-rich nature (10.9% crude fat, based on manufacturer’s information), which translates into richer dietary calories [54]. This hypothesis is supported by high expression of the fat-digesting enzyme, pancreatic triacylglycerol lipase (PNLIP) (Fig. 3G and Additional file 7: Table S2), and overrepresentation of fatty acid degradation processes in FF-fed snails (Figs. 4 A and D). Considering previous proximate composition analysis of the effects of drying on leafy vegetables [55], our DL diet may have higher energy value than FL or WL, hence its better performance on snail growth and reproduction.

Higher survival in snails fed on FF and SG may be partly due to the upregulated expression of many innate immune protein-coding genes and enrichment of immune-related pathways, such as glycoprotein and glycosphingolipid synthesis. These genes included BPI/LBP, serine protease inhibitors (e.g. SERPINs), MUC5AC, APLY-A, apextrin C (APEC), and C-type lectins including perlucin (PLC)-like protein, mannose receptor 1 (MRC1), lectin ADEL, ladderlectin, and biomphalysins (2, 4, 7, 10, 20 and 21) (Additional file 6: Dataset 3). Since laboratory snail maintenance conditions are usually not aseptic and may support rapid growth of harmful microorganisms [10], these immune genes may fortify snail immunity against such microbes, thereby enhancing snail fitness for survival. For example, we found that increased immune gene expression correlated with reduced presence of *Chlamydiia* bacteria, which are predominantly obligate intracellular pathogens (Fig. 6F).

In contrast, FL snails exhibited high transcriptional activation of xenobiotic detoxification pathways (Figs. 4 C and D), suggesting potential exposure to dietary toxins. In these snails, we noted upregulation of cytochrome P450 (CYP450) enzymes and glutathione transferases (GSTs), which mediate sequential biotransformation and conjugation of xenobiotics for excretion [56, 57]. Other xenobiotic-metabolising enzymes including FMN-dependent NADPH-azoreductase (AZR) [58], dye-decolorising peroxidase YFEX [59], xanthine dehydrogenase (XDH) [60], zeta-crystallin (CRYZ) [61], UDP-glucuronosyltransferase 2A3 (UGT2A3) and sulfotransferases (SULT1A1 and SULT1B1) [62] were also upregulated in FL (Additional file 6: Dataset 3; Additional file 7: Table S2), suggesting the increased need for FL snails to counter diet-related toxins. Also, transcriptional activation of nuclear receptors (NRs) and cognate hormone ligands and cytokines (predominantly IL17 based on our data) (Figs. 4C, E and F) is instrumental in mounting cellular xenobiotic response [63], while increased expression of genes associated with caveolae (cup-shaped uncoated invaginations at the plasma membrane) (Fig. 4C) is indicative of acute mechanical stress [64]. Concurrently, some transcriptionally elevated enzymes such as pirin (PIR) and oxidoreductase HTATIP2 (Figs. 3G and 3H) are markers of oxidative stress and apoptosis [65, 66]. Xenobiotics in water lettuce likely originated from bioaccumulation of environmental pollutants, such as microcystins (cyanogenic toxins) from agal bloom, which were recently detected in Lough Neagh, the UK and Ireland’s largest lake [67]. Additionally, some inherent compounds in green leafy vegetables, including nitrates, phytates, tannins, oxalates, and cyanogenic glycosides, are involved in plant defence and can induce cellular damage in invertebrates when heavily consumed [68]. Similarly, quinones, which are produced in lettuce postharvest [69], are cytotoxic and can cause oxidative damage or even death in animals [70].

Growth factors and regulators are essential for molluscan growth and development [71]. We identified four molluscan insulin-related peptide 3 (MIP3)-like genes (MIP3.1, MIP3.2, MIP3.3 and MIP3.4) and an insulin-like growth factor-binding protein (IGFBP-rP1). These were however upregulated in the stunted FL-fed snails, except MIP3.4 that showed upregulation in FF and SG snails (FF log2fc = 4.63, *p*adj = 2.61e-09; SG log2fc = 4.00, *p*adj = 1.79e-06) (Additional file 6: Dataset 3). Other growth-related genes encoding perlucin, temptin, and dermatopontin proteins that were highly expressed in FF and SG play important roles in promoting shell biomineralisation and shell size in molluscs [72–74]. Additionally, functional enrichments displayed in Figs. 4A and 4B suggest conserved orchestration of mitotic cell proliferation in FF and SG snails. For example, minus-end-directed microtubule activity mediates spindle fibre morphogenesis [75], dynein intermediate chain binding facilitates microtubule organisation and centrosome replication and separation at interphase [76], while dynein light intermediate chain binding is required for mitotic spindle positioning [77]. Other gene factors mediating growth activities are listed in Figs. 4E and 6E. Altogether, these data highlight the network of molecular triggers underpinning diet-responsive growth observed with the high-caloric diets.

Reproduction in *B. glabrata*, as with other molluscs, is governed largely by peptidergic neuroendocrine signalling [78] as well as non-endocrine gene regulation [79]. Among the most significantly expressed non-endocrine genes in snails treated with high-caloric diets in this study were BPI/LBP, HCY1, and TEMPT (Figs. 3G and H; Additional file 7: Table S2). These genes play multifunctional roles including reproduction in *B. glabrata*. In other studies, experimental knockdown of the BPI gene resulted in a significant reduction of oviposition in *B. glabrata* [80], while HCY1 expression is reported to directly correlate with the snail sexual maturity [81]. Although TEMPT may not directly regulate egg-laying in *B. glabrata*, it serves as chemosensory attractant among conspecifics [82], therefore promoting copulatory behaviour that may enhance reproductive success.

Angiotensin-converting enzyme (ACE, a dipeptidyl carboxypeptidase) and two angiotensin receptors, ATRAP and AT1B, were upregulated in FF and SG snails (Figs. 3G and 3H, S4 Table). ACE processes cleaved inactive angiotensin I into bioactive angiotensin II [83], a peptide hormone that is associated with fertilisation in invertebrates, including *Bombyx mori* (insect) [84] and *Crassostrea gigas* (mollusc) [85]. Upregulation of this gene may play similar role in *B. glabrata.* Our transcriptional analysis pipeline also identified seven differentially expressed FMRFamide receptor-like genes, one of which (LOC106063694) was upregulated in SG, while the other six were downregulated. In FF, two (LOC129924834 and LOC106063694) were upregulated and five downregulated (Additional file 6: Dataset 3). These genes encode neuropeptides that contain a C*-*terminal Arg-Phe-NH_2_ (RFamide) motif which are essential regulators of molluscan reproduction [86]. For example in *B. glabrata*, FMRFamide induces preputium eversion in early copulation [87]. Other reproduction-related DEGs, such as spermatogenesis-associated protein 7 (SPATA7) are listed in Fig. 6G.

Higher reproductive capacity in FF vs SG snails is reflected in altered expression of reproduction-related genes. Three VLDLR-like genes and four LDLR-like genes were more highly expressed in FF than SG snails (Fig. 3I) (Additional file 6: Dataset 3). These two lipoprotein receptors are involved in uptake of yolk constituents into the oocyte, as documented in a wide range of metazoans including nematodes [88], insects [89], gastropods [90], crustaceans [91], and fishes [92].

The effect of diet on *B. glabrata*’s gut microbiome was reported by a previous publication [93]. We build on that study by examining the effect of diet on whole-snail microbiome in the gut and other organs, using metatranscriptomic tools. Our initial KEGG analysis highlighted beneficial nutritional signatures of gut microbiota through enrichment of butanoate and propanoate pathways in FF snails. Butanoate and propanoate are primarily short-chain fatty acid (SCFA) metabolites of gut microbial saccharolytic, and to a minor extent, proteolytic, fermentation [94], which inhibit growth of harmful bacteria, facilitate gut nutrient absorption, maintain homeostasis, and reduce inflammation and oxidative stress [95, 96]. *Bacilli* and *Clostridia* of the phylum *Firmicutes* are major producers of SCFAs [96], with certain species used as probiotics to promote health, growth and reproduction in fish and shellfish aquaculture [95]. Nonetheless, members of *Actinomycetes*, *Spirochaetia*, *Thermotogae*, and *Fusobacteriia* also contribute significantly to SCFA production [97], all of which were more transcriptionally abundant in FF and SG snails in line with the observed phenotypes in these groups. Gene–taxon correlation analysis (Fig. 6E–G), however, did not support the beneficial relevance of *Bacilli*, *Clostridia*, and *Actinomycetes*, despite these being the top relatively abundant taxa (Fig. 5E). This may have been due to compositional expansion of less beneficial species within these taxa. While FF- and SG-enriched classes such as *Epsilonproteobacteria*, *Terriglobia*, *Desulfobulbia*, *Desulfobacteria*, and *Gemmatimonadetes* are not SCFA producers, they can be found in the gut and other organs, benefitting their aquatic invertebrate hosts via degradation of complex polysaccharides, fatty acids and proteins, and hydrogen sulfide detoxification [98–101]. These data, coupled with our Spearman correlation analysis, support dietary alteration of snail microbiomes towards enhanced growth, immunity and reproductive output in SG and FF snails compared to FL snails.

Indicator taxa found in the diet groups may potentially serve as ecological indicators of dietary exposures or changes. While the two anaerobic indicator taxa associated with FF and SG diets are poorly understood, we do know that *Candidatus Cloacimonadia* can be found in lipid-rich environments where they perform nutrient cycling, generating acetate (comprising 60% of total SCFAs [96]) from carbohydrate fermentation or propionate oxidation [102, 103]. On the other hand, *Candidatus Brocadiia* are known for anaerobic ammonium oxidation, removing ammonium from aquacultural systems to reduce nitrogen pollution and enhance aquacultural health [104]. Taken collectively, FF and SG diets altered snail microbiomes and established consortia of functionally useful microbes, with wide-ranging benefits to the snails, correlating with enhanced reproduction, growth and survival.

## Limitations

In this study, snail samples were collected for sequencing only at a single timepoint. Our RNA-Seq data is therefore age-specific and may only reflect (meta)transcriptomic landscapes around eight weeks post-feeding. Additionally, the NMRI strain of *B. glabrata* that we used has been maintained under laboratory conditions for many years, and as such may not be representative of other lab strains or wild snails, especially in terms of snail exposure to more diverse food sources, microbes, parasitic infections and other confounding environmental variables.

## Conclusions

This study reports new data that significantly enhances understanding of *B. glabrata* diet– gene–microbiome–phenotype interactions. Laboratory feeding of *Biomphalaria* snails with a single lettuce diet is common because it represents a low cost, accessible food source. We have shown that its low-calorie composition, potentially harmful phytochemicals, potential contamination with environmental toxins, and non-prebiotic nature mark it as a less-than-ideal food source for maintenance of *B. glabrata*. We have highlighted that a simple alternative food source, commercially available fish food flakes, can significantly enhance *B. glabrata* fecundity and survival. This, we hope, will inform dietary precision and optimisation in laboratory maintenance and enhance large-scale rearing of *Biomphalaria* snails that can bolster laboratory culture of the *S. mansoni* life cycle. We provide key mechanistic insights into the molecular and microbial basis of these phenotypes including a wealth of putative genes and pathways that govern key physiological processes in the snail. Functional genomics approaches should focus on characterising these genes and extending similar studies to wild type snail isolates to uncover new potential targets for molecular snail control.

## Supplementary information

**Additional file 1: Fig. S1. Global heatmap for *Biomphalaria glabrata* gene expression across all samples based on dietary feeding.**

The colour gradient scale represents the Z-score (scaled) expression of a gene in a sample. Positive values indicate the gene expression in the sample is above the mean expression of the same gene across all samples. Negative Z-score indicates that the sample has lower expression for that gene compared to the gene’s average expression across all samples. At zero Z-score, expression of the gene in the sample equals to the average expression of that gene across all samples. Therefore, the global heatmap only reflects relative gene expression patterns across conditions, not statistically significant up- or down-regulation between groups.

**Additional file 2: Fig. S2. Microbiome library size per sample.** This was estimated based on raw operational taxonomic units (OTUs) and represents the total number of RNA sequence reads generated for a single biological sample.

**Additional file 3: Table S1. Read quality and mapping data per sample.**

Sheet 1 presents the numerical summary of the RNA-Seq data pre-and post-trimming. Column A – sample file name; Column B – sample symbol; Column C – number of paired-end reads pre-trimming; Column D – number of reads post-trimming; Column E – number of base pairs processed; Column F – number of base pairs that passed quality filtering; Column G – percentage of base pairs kept post-filtering; Column H – number of unpaired reads after quality filtering; Column I – percentage of reads with adapters; Column J – percentage of base pairs trimmed; Column L – average read length; Column M – percentage GC content of the reads. Sheet 2 presents the mapping statistics of the reads per sample. Column A – sample file name; Column B – sample symbol; Column C – number of mappable reads; Column D – number of reads that mapped with the *Biomphalaria glabrata* reference genome; Column E – number of unmapped reads; Column F – percentage of mapped reads; Column G – percentage of unmapped reads; Column H – percentage of unmapped reads that was classified taxonomically by Kraken 2.

**Additional file 4: Dataset 1. Gene count matrix.**

Gene assembly and counts from StringTie.

**Additional file 5: Dataset 2. DESeq2 filtered gene counts.**

Gene counts after the removal of lowly-expressed genes.

**Additional file 6: Dataset 3. Metadata on all differentially expressed genes (DEGs) in the three pairwise comparisons with annotations.**

DESeq2 result data of FL vs FF (Sheet 1), FL vs SG (Sheet 2), and SG vs FF (Sheet 3) with gene names and symbols. Only DEGs with log2fc above or below 1.0 and *p*adj < 0.01 were included.

**Additional file 7: Table S2. List of top 20 upregulated and top 20 downregulated differentially expressed genes (DEGs) in pairwise comparisons.**

FL vs FF (Sheet 1), FL vs SG (Sheet 2), SG vs FF (Sheet 3). Tables include chromosome number and number of exonic sites for each protein-coding gene, curated from NCBI Batch Entrez. Top genes were ranked based on significance *p*adj level. NA* represents ‘Not Applicable’.

**Additional file 8: Dataset 4. Upregulated gene ontology (GO) enrichments based on diet.** This table consists of the upregulated GO terms (biological processes BP, cellular components CC, and molecular functions MF) in snails fed on fish food (FF), snail gel (SG) and fresh lettuce (FL) diets. For each GO term, ShinyGO enrichment FDR values (FDR ≤ 0.05), number pathway genes (nGenes) found differentially expressed, number of pathway genes in our *Biomphalaria glabrata* background gene list, fold enrichment values, and list of genes associated with the pathway were presented.

**Additional file 9: Dataset 5. Upregulated Kyoto Encyclopaedia of Genes and Genomes (KEGG) pathways based on diet.**

This table consists of the upregulated KEGG pathways in snails fed on fish food (FF), snail gel (SG) and fresh lettuce (FL) diets. For each KEGG pathway, ShinyGO enrichment FDR values, number pathway genes (nGenes) found differentially expressed, number of pathway genes in our *Biomphalaria glabrata* background gene list, fold enrichment values, and list of genes associated with the pathway were presented.

**Additional file 10: Dataset 6. Raw operational taxonomic unit (OTU) data.**

Bracken output after taxonomical denoising. Taxa printed in pink colour indicate yeasts, red colour indicates viruses, green colour indicates protists, blue colour indicates archaea, and black colour indicates bacteria.

**Additional file 11: Table S3. Statistical data on microbial relative abundance.** Kruskal–Wallis significance result (Sheet 1) and pairwise Mann–Whitney *U* significance result. *p*-values were FDR-adjusted (Sheet 2).

**Additional file 12: Dataset 7. Analysis of compositions of microbiomes with bias correction (ANCOM-BC) result data.**

In the result data, lfc_(intercept) indicates the log fold-change of the baseline or reference FL group. lfc_DietFF indicates the estimated log fold-change of FF relative to the baseline FL group. Positive values mean higher in FF and negative values mean lower in FF. lfc_DietSG indicates the estimated log fold-change of SG relative to the baseline FL group. Positive values mean higher in SG and negative values mean lower in SG. se_(Intercept) shows the standard error of the intercept (uncertainty around the baseline group estimate). Standard error of the FF log fold-change (se_DietFF) and of the SG log fold-change (se_DietSG); Wald test statistics (estimate divided by standard error) for the intercept (W_(Intercept)), for the FF log fold-change (W_DietFF), and for the SG log fold-change (W_DietSG); Raw *p*-value for the intercept (p_(Intercept)), for the FL_vs_FF test (p_DietFF), and for the FL_vs_SG test (p_DietSG); FDR-corrected *p*-value for the intercept ((q_(Intercept)), for the FL_vs_FF test (q_DietFF), and for the FL_vs_SG test (q_DietSG), were included. diff_DietFF is ‘TRUE’ if FF was differentially abundant relative to FL, and diff_DietSG is ‘TRUE’ if SG was differentially abundant relative to FL. passed_ss is ‘TRUE’ if the taxon passed the structural zero check for that diet group. lfc, se, W, *p*, and *q* values for SG_vs_FF indicate the estimated log fold-change, standard error, Wald test, raw *p*-values, and FDR-corrected *p*-values of the inter-diet comparison, respectively.

**Additional file 13: Text S1. Indicator taxa result summary.**

In this multilevel pattern analysis, IndVal stat represents how strongly a taxon indicates a group, A (specificity) indicates the taxon exclusivity to the group, B (fidelity) indicates the degree of consistency of the taxa in that group, and *p*-value shows the significance of the association.

## Abbreviations

ANCOM-BC: Analysis of compositions of microbiomes with bias correction
CLR: Centred log-ratio
DA: Differential abundance
DAVID: Database for annotation, visualization, and integrated discovery
DL: Dried lettuce
FF: Fish food
FL: Fresh lettuce
GO: Gene ontology
HMMER: Hidden Markov model
KEGG: Kyoto encyclopaedia of genes and genomes
OTU: Operational taxonomic unit
RA: Relative abundance
SCFA: Short-chain fatty acid
SG: Snail gel
WL: Wilted lettuce.

## Acknowledgements

We are thankful to Dr Gabriel Rinaldi for generously providing the initial population of NMRI *Biomphalaria glabrata*.

## Declarations

### Availability of data and materials

Raw sequencing data have been deposited in the European Nucleotide Archive (ENA) (https://www.ebi.ac.uk/ena) under accession number PRJEB96928. All other relevant data and metadata are provided within this manuscript and in the supplementary files. All R scripts, Linux bash scripts, and some associated data entry files used in this study are accessible on Zenodo repository at https://doi.org/10.5281/zenodo.17058355.

### Authors’ contributions

DOF: Conceptualization, Data curation, Formal analysis, Investigation, Methodology, Project administration, Software, Visualization, Writing – original draft, Writing – review & editing. CL: Conceptualization, Data curation, Investigation, Methodology, Writing – review & editing. DW: Conceptualization, Data curation, Methodology, Software, Writing – review & editing. GNG: Conceptualization, Methodology, Supervision, Validation, Writing – review & editing. PM: Conceptualization, Funding acquisition, Methodology, Project administration, Resources, Supervision, Validation, Writing – review & editing. All authors read and approved the final version for submission.

