## Supplementary figures and images for "Dietary variations drive divergent phenotypic, transcriptomic, and metatranscriptomic profiles in *Biomphalaria glabrata*, a schistosomiasis vector snail"

### Additional file 1 and 2_Suppl. figures

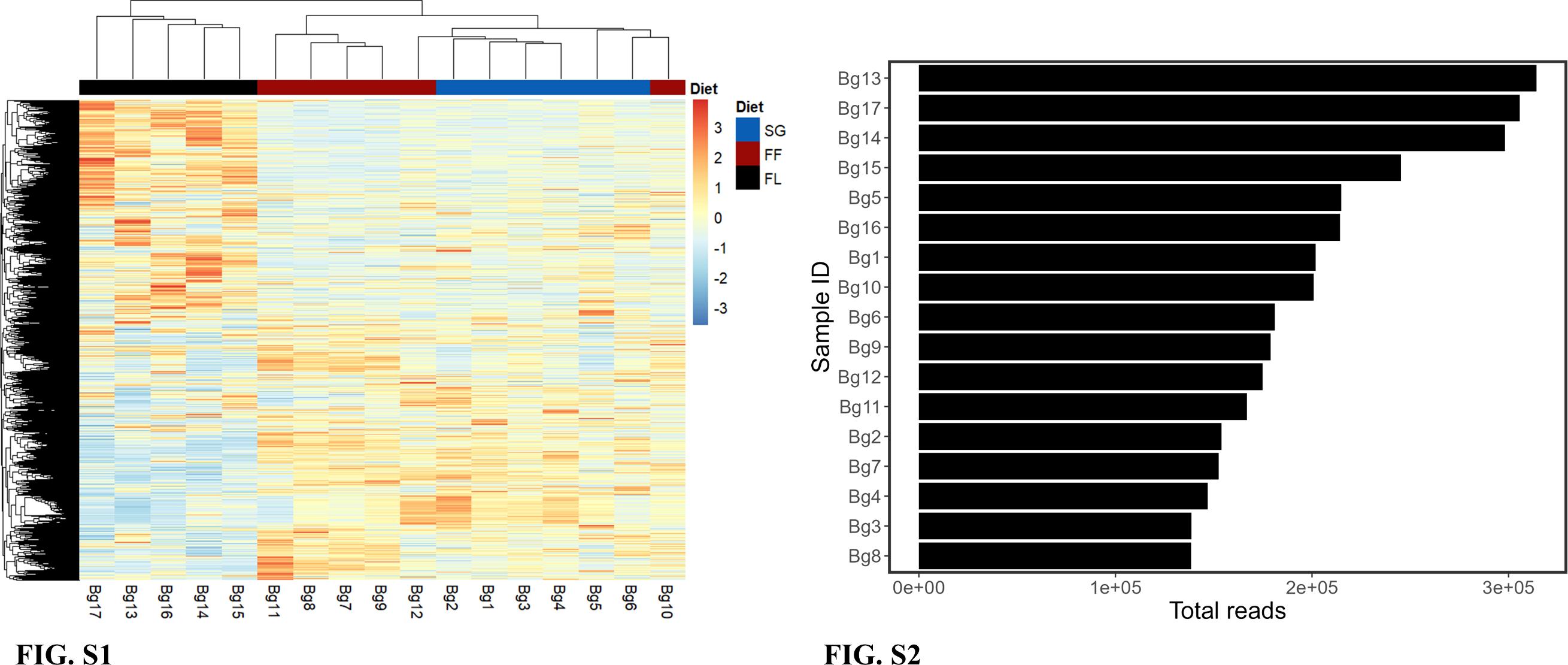
